# Hemodynamic and Microvascular Adaptations to Aerobic Training Intensity Improve Maximal Oxygen Consumption

**DOI:** 10.1101/2025.11.17.688970

**Authors:** E. Maufroy, C. Rigaut, C. Maufroy, N. Baeyens, G. Deboeck

**Affiliations:** Research Unit in Rehabilitation Sciences, Faculty of Human Movement Sciences, Université libre de Bruxelles (ULB), Brussels, Belgium; Laboratoire de Physiologie et Pharmacologie, Faculty of Medicine, Université ibre de Bruxelles (ULB), Brussels, Belgium; Transferts, Interfaces and Processes, Ecole Polytechnique de Bruxelles, Université Libre de Bruxelles (ULB), Brussels, Belgium

## Abstract

**Background:** Aerobic training enhances VO₂_max_, yet the contribution of peripheral microvascular remodeling to this improvement remains insufficiently understood. This research demonstrates how two distinct training modalities, high-intensity interval training (HIIT) and moderate-intensity continuous training (MICT), influence oxygen transport dynamics and microvascular remodeling.

**Methods:** Twenty-five healthy sedentary adults (15 women, 10 men; mean age 25 ± 2 years; normal BMI) were randomly assigned to HIIT or MICT for 8 weeks. VO₂_max_ was assessed before and after the training program. 15 participants underwent non-invasive maximal cardiac output measurement (Q_max_), while vastus lateralis muscle biopsies were obtained from 10 participants. Tissue samples were cleared and immunolabeled for VE-cadherin and alpha-smooth muscle actin to observe microvasculature architecture. A computational hemodynamic model integrating cardiac output and microvascular parameters was constructed to estimate flow dynamics.

**Results:** VO₂_max_ increased significantly in both training groups, with a greater improvement in HIIT (p = 0.024). Q_max_ increased similarly in both groups (p = 0.001), while calculated arteriovenous oxygen difference (a-vO₂ diff) showed a trend toward improvement only in HIIT. No formation of new capillaries nor anastomoses (angiogenesis) was detected in either group; however, both HIIT and MICT induced significant capillary and venule dilation. Notably, only HIIT led to a significant increase in pericyte coverage (p = 0.047). Venules of both groups exhibited dilation accompanied by increased surrounding smooth muscle cells. No remodeling was found in arterioles. Hemodynamic modelisation estimated higher shear stress during HIIT than MICT and vasodilation tended to decrease shear stress over time during both training. Furthermore, pericyte recruitment was modelized to adapt to shear stress level limiting excessive capillary dilation during high effort intensity.

**Conclusion:** HIIT induces superior improvements in VO₂_max_ and distinct microvascular structural adaptations rather than angiogenesis. HIIT is supposed to stimulate a protective adaptation at the capillary level, limiting excessive dilation during maximal effort. Our hemodynamic model supports this shear stress-dependent mechanism. These findings underscore the role of exercise intensity and hemodynamics in shaping microvascular responses to endurance training.

**Clinical Perspective:**

- Peripheral adaptation to exercise is linked with the dilation of muscle capillaries and venules.
- Mechanoadaptive responses, rather than growth factor-mediated angiogenesis, drive the remodeling of the muscle microvasculature.
- High-intensity interval training elicits higher shear stress than moderate continuous interval training, linking the adaptation of the microvasculature to increased blood flow as the primary factor that explains the superiority of HIIT compared to MICT in improving maximal oxygen consumption.

**Clinical implication:**

- Training regimens should focus on increasing peripheral flow and shear stress to initiate microvasculature remodeling.
- Potentiating mechanoadaptative responses and microcirculation remodeling would provide a means to improve cardiovascular function and fitness

## Introduction

Aerobic exercise capacity, measured by maximal oxygen consumption (VO_2max_) is directly related to overall health, notably to low risk of cardiovascular disease and all-cause mortality (1). Improving VO_2max_ requires regular exercise training, which induces repeated disruptions of homeostasis that promote beneficial physiological adaptations (2). In this context, exercise training modalities such as moderate-intensity continuous training (MICT) and high-intensity interval training (HIIT) (3) have been successfully used for decades (4). MICT is typically performed over a relatively extended period of time between 50% and 70% VO₂_max_, corresponding to efforts below or near the first ventilatory threshold (VT1). VT1 reflects an individual’s aerobic capacity and marks a shift in substrate utilization, transitioning from predominantly aerobic pathways to increased engagement of anaerobic metabolism. HIIT, on the other hand, alternates bouts of high-intensity exercise (above the VT1 threshold) with recovery phases. Ongoing debate, however, persists as to whether HIIT yields similar or superior improvements in VO_2max_ compared to MICT (5–7) and over the exact mechanism of action. The Fick principle states that VO_2_ is defined as the product of cardiac output (Q) and the arteriovenous oxygen difference (a-v O₂ diff). Improvement in aerobic capacity is therefore subordinated to central enhancements (e.g., increased cardiac contractility, stroke volume) and peripheral enhancements (e.g., improved aerobic metabolism, enhanced muscle tissue perfusion) (8,9). Increased exercise intensity leads to a proportional rise in hemodynamic forces within the microcirculation (arterioles, capillaries, and venules) of active muscles. Hemodynamic forces, represented by fluid shear stress (FSS) and vessel wall circumferential stretch (10) (11–13), are sensed by mechanoreceptors located between endothelial cells and vascular smooth muscle cells, triggering intracellular signalling cascades possibly driving vascular remodelling (14–16). The sequential nature and elevated intensities of HIIT may provoke higher, more dynamic and repeated hemodynamic stimuli, potentially leading to more pronounced vascular remodelling. This study aims to elucidate how peripheral adaptations, particularly microvascular adaptive remodelling, contribute to improved cardiovascular efficiency and maximal oxygen uptake (VO₂_max_) following different exercise regimens. By comparing HIIT and MICT, we seek to characterize intensity-dependent vascular adaptations and their role in optimizing tissue perfusion, oxygen delivery, and aerobic capacity.

## Methods

### Study design

This prospective study was conducted between September 2022 and December 2024 within the Research Unit of Rehabilitation Sciences at Université libre de Bruxelles. All participants received detailed information about the study and provided written informed consent, which was approved by the Research Ethics Committee of the Brussels University Hospital (B4062021000227). Inclusion criteria required participants to be between 18 and 40 years old, to have a body mass index (BMI) below 30, and be classified as sedentary. Sedentary status was defined as self-reported physical activity levels below the World Health Organization guidelines, less than 150 minutes of moderate-intensity or 75 minutes of vigorous-intensity exercise per week (1). Exclusion criteria included smoking, regular medication use, and any history of cardiovascular, respiratory, or metabolic diseases. The study protocol is illustrated in *Figure 1*. Twenty-five participants were recruited and underwent a cardiopulmonary exercise test (CPET) to determine their maximal oxygen uptake (VO₂_max_). Of those 25 subjects, a subgroup of 15 participants completed a second exercise test within 24 hours and following a similar workload increment, with measurement of non-invasive cardiac output at rest and at maximal exercise workload (Q_max_). VO₂_max_ and Q_max_ were subsequently used to derive the maximal arteriovenous difference (a-v O₂ diff max) according to the Fick principle. Another subgroup of 10 participants underwent a resting muscle biopsy of the vastus lateralis (quadriceps). Muscle samples were collected, fixed and processed as detailed below (*Muscle Biopsies* section). All participants were then enrolled in an 8-week training program, 3 times a week, and randomly assigned to either a MICT group or HIIT group. At the end of the 8-week intervention, all baseline assessments were repeated. Cardiac output measurements and muscle biopsy data were used to construct a hemodynamic model (*Hemodynamic model* section) designed to link macro-level cardiovascular adaptations with micro-level vascular changes.

**Figure 1:**
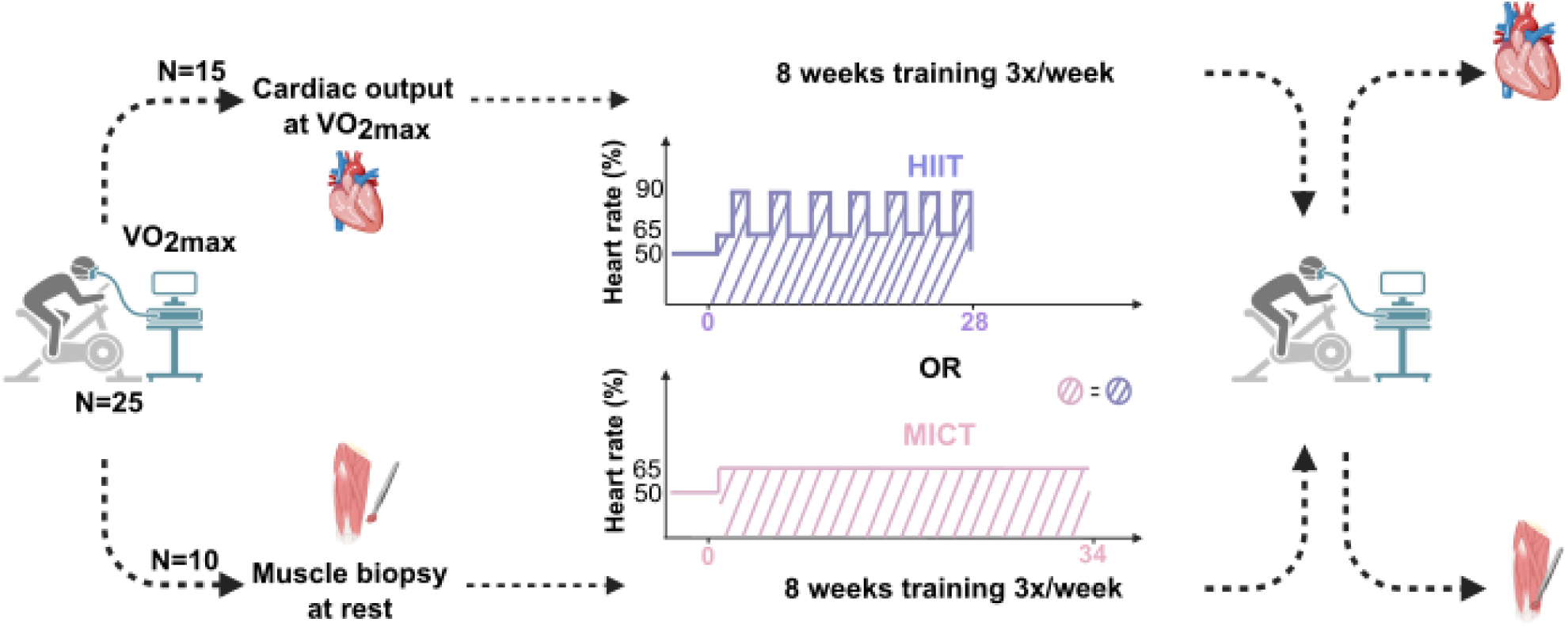
Schematic overview of the study protocol and assessment timeline. Maximal oxygen uptake (VO₂_max_) at baseline and following an 8-week training intervention. A subset of 15 participants underwent cardiac output measurements at VO₂_max_, while another subset of 10 participants underwent muscle biopsy of the vastus lateralis at rest. Participants were randomly assigned to either a high-intensity interval training (HIIT) or moderate-intensity continuous training (MICT) group. Both training modalities were matched for hemodynamic load, defined as the product of session duration and target intensity, individualized according to each participant’s percentage of maximal heart rate.

### Training modalities

Cycling-based exercise training was performed under supervision three times per week. Throughout each session, heart rate was continuously monitored with visual feedback to facilitate adherence to prescribed intensity targets. The training workload was adjusted every five sessions, following individualized heart rate zones. Both exercise protocols were designed to elicit an equivalent hemodynamic load, defined as the product of training duration and targeted heart rate. All training sessions (HIIT and MICT) commenced with a standardized 3-minute warm-up at 50% HR_max_. The 28-minute HIIT protocol consisted of seven 2-minute bouts at 90% HR_max_, each separated by 2 minutes of moderate-intensity exercise at 65% HR_max_. The 34-minute MICT protocol involved continuous exercise at 65% HR_max_ for 34 minutes. HR_max_ was determined from the initial cardiopulmonary exercise test (CPET).

### Cardiopulmonary exercise test

CPET was conducted on a cycle ergometer (Ergoselect 100, Ergoline GmbH, Germany) using a one-minute workload increment protocol. Participants were instructed to maintain a pedaling cadence between 60 and 70 revolutions per minute throughout the test. Oxygen consumption (VO₂), carbon dioxide production (VCO₂), and ventilation (VE) were measured breath-by-breath via a face mask (COSMED, Rome, Italy). VO₂_max_ was defined as the highest VO₂ value recorded over a 20-second interval at peak exercise. Additional methodological details, including test initiation procedures, criteria for maximal effort, and determination of the first ventilatory threshold, are provided in the Supplemental Methods (online appendix).

### Non invasive measure of cardiac output

Cardiac output (Q) was measured non-invasively using the inert gas rebreathing method with the Innocor device (COSMED, Rome, Italy), which operates based on a single-alveolar lung model. Stroke volume was calculated by dividing the cardiac output (Q) by the heart rate (HR). Participants rebreathed a gas mixture containing 0.5% nitrous oxide (NO, blood-soluble) and 0.1% sulfur hexafluoride (SF₆, blood-insoluble) diluted with ambient air. Pulmonary shunting is minimal in healthy individuals; pulmonary blood flow (PBF) was assumed to be equivalent to Q (PBF ≈ Q). Additional technical details are provided in the Supplemental Methods (online appendix).

### Muscle Biopsies

Resting muscle biopsies were obtained from the vastus lateralis of the quadriceps 14 days before the baseline CPET and 5 days after the post-training CPET to minimize any discomfort or pain that could interfere with exercise performance. To avoid sampling scar tissue and ensure anatomical consistency, the first biopsy was performed on the right leg and the second on the left. Muscle samples were processed using the PEGASUS protocol, a modified tissue-clearing technique optimized for immunolabeling and three-dimensional imaging. VE-cadherin and α-smooth muscle actin (α-SMA) were used to label endothelial borders and vascular smooth muscle structures, respectively, allowing for the precise delineation of the intima and media layers. Confocal imaging was performed using a Nikon AX R confocal microscope. For each sample, z-stack acquisitions at 10× magnification were stitched together to generate wide-field composites that encompassed the full tissue volume. From these representative regions, areas containing arteriolar, capillary, and venular segments were selected for high-resolution imaging at 20× magnification. Vessel diameters were quantified transversely using NIS-Elements software, based on fluorescence-defined boundaries. Intima diameters were defined by VE-cadherin signal, while media diameters corresponded to α-SMA-positive structures. Measurements were performed independently on each fluorescence channel to ensure layer-specific morphometric accuracy. Capillary density was assessed in two anatomically distinct regions per sample using a volumetric approach that combined z-stack depth with a fixed surface area of 200 × 200 μm, with a mean imaging depth of 70 ± 6 μm at baseline and 83 ± 23 μm following the training intervention. Additional procedural details regarding biopsy collection, tissue preparation, immunostaining, and clearing steps are provided in the Supplemental Methods (online appendix).

### Hemodynamic model

The arterial side of the vascular network supporting the hemodynamic model consisted of five dichotomous branches of the arteriole, resulting in 32 capillaries per arteriole. The venous side of the network consisted of five dual merges, giving one venule per arteriolar *(Figure 4, panel A)*.

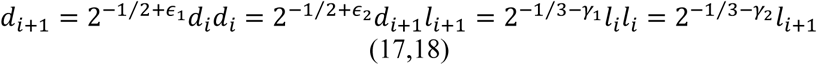

**Figure 2.**
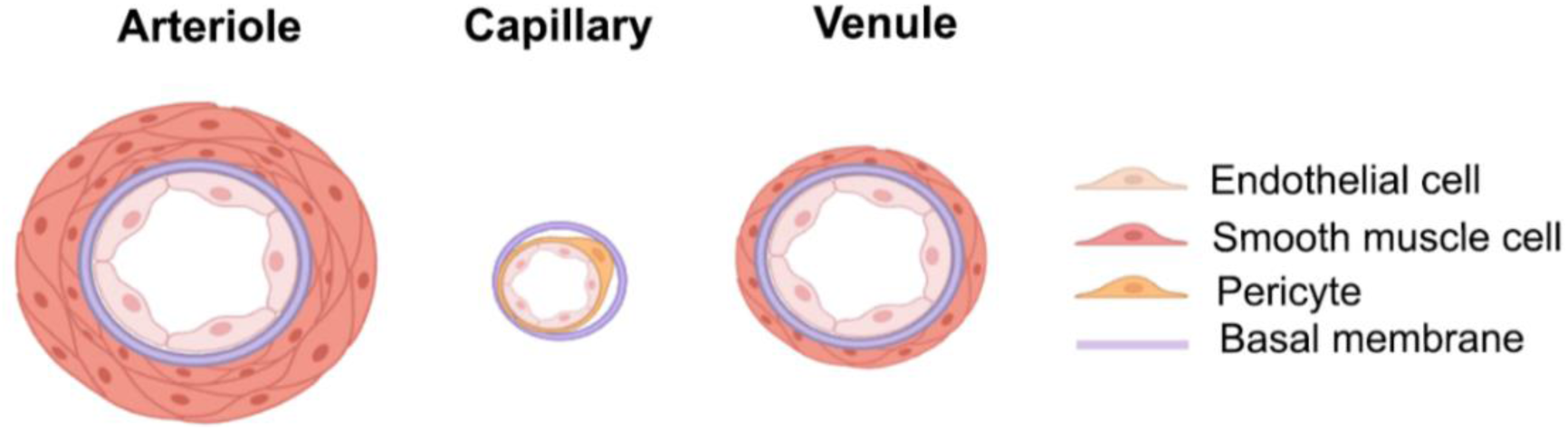
Schematic representation of the microvascular wall architecture, used as the structural basis for the development of the hemodynamic model.

**Figure 3:**
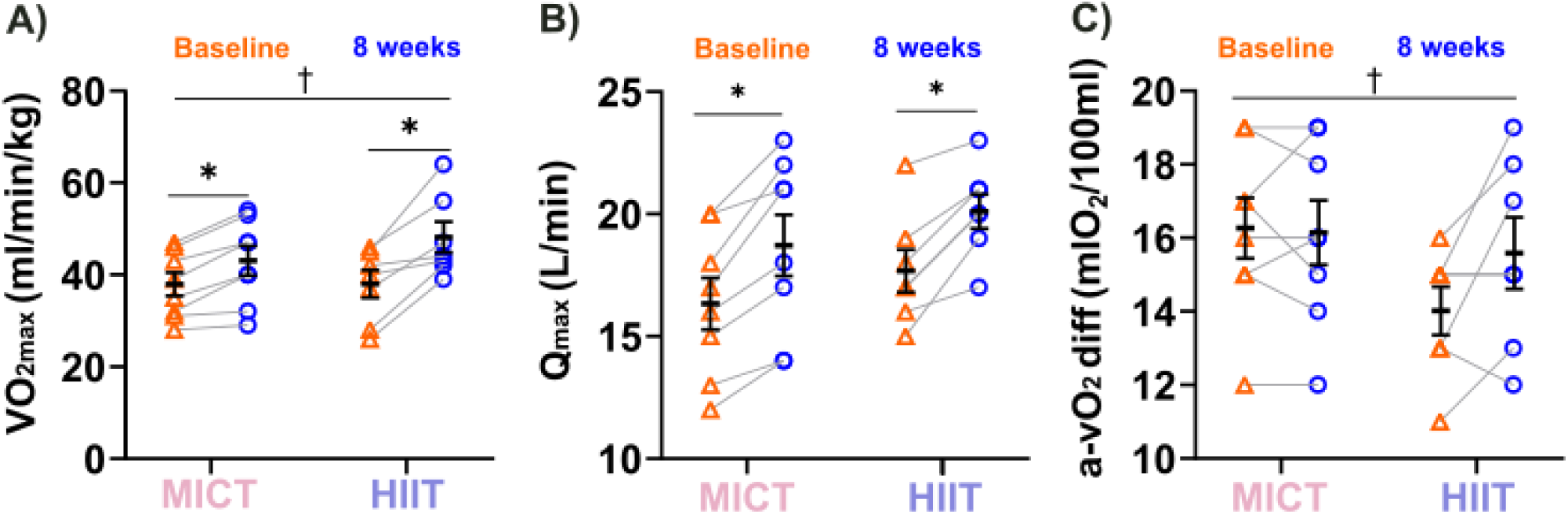
Cardiac output and oxygen transport adaptations following 8 weeks of MICT or HIIT (N=15). (A) illustrates the evolution of VO₂_max_ across the training period. Both MICT and HIIT groups demonstrated significant improvements post-intervention; however, the magnitude of increase was more pronounced in the HIIT cohort. (B) depicts changes in cardiac output measured at VO₂_max_. Both training modalities elicited comparable increases in peak cardiac output. (C) presents the maximal arteriovenous oxygen difference (a-vO₂ diff), calculated using the Fick principle (VO₂/Q). The HIIT group demonstrated a trend toward greater enhancement in a-vO₂ diff compared to MICT, with a distinct temporal evolution between groups. Orange triangles represent baseline data; blue circles indicate post-training. **Statistics:** Normality tests were tested with a Shapiro-Wilk test. ANOVA tests were performed to compare groups, when significant, Tukey post hoc tests were applied. For pairwise comparisons, Student’s t-tests were used. Results are presented as mean ± SD. * denotes a significant difference between pre– and post-training; † indicates a significant difference between time*groups.

**Figure 4:**
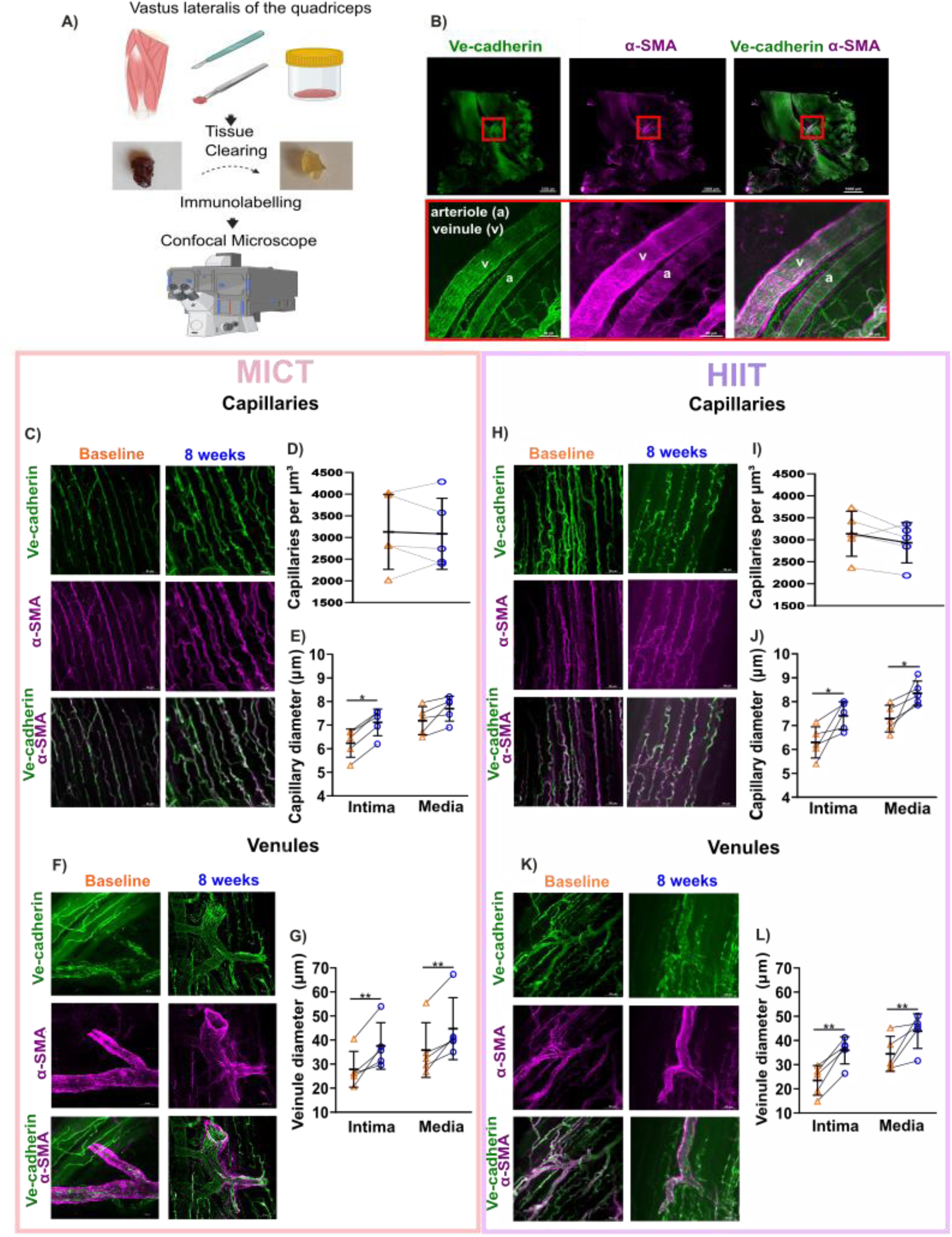
This figure presents the methodological workflow for muscle biopsy processing and image acquisition, as well as the vascular remodeling observed in response to exercise training. (A) Schematic overview of the biopsy protocol performed in 10 participants. Samples were collected under local anesthesia from the vastus lateralis using a scalpel, then cleared and immunolabeled with VE-cadherin to visualize endothelial cells and withd α-smooth muscle actin (α-SMA) to identify smooth muscle cells and pericytes. Image acquisition was performed using a Nikon AX R confocal microscope. (B) Overview of an entire muscle sample (scale bar: 1000 µm; objective: Apo 10×), with the red box indicating the region of interest. This region contains a venule and an arteriole, distinguishable by their structural features: venules exhibit a loose endothelial architecture with disorganized smooth muscle coverage, whereas arterioles display tightly arranged endothelial cells and concentric smooth muscle rings (scale bar: 50 µm; objective: Apo 20×). (C-G) correspond to the MICT group and (H-L) to the HIIT group. (C and H) represent the visualization of the capillary network before and after 8 weeks of training while (F et H)represent the visualization of a venule before and after 8 weeks of training (scale bar: 50 µm; objective: Apo 20×). (D and I) is the quantification of capillary network, both group didn’t succeed to increased the number of capillaries after training. (E and J) quantify capillary diameter changes at baseline and 8 weeks after training, measured at the intima (VE-cadherin) and media (α-SMA) levels. Both groups exhibited significant dilation of the capillary intimal diameter following training; however, only the HIIT group (panel J) demonstrated a concomitant increase in pericyte coverage. (G and L) demonstrate venular remodelling in both groups, with increased intima diameter and enhanced smooth muscle cell coverage. Orange triangles represent baseline data; blue circles indicate post-training. **Statistics:** Normality tests were performed by Shapiro-Wilk test. For pairwise comparisons, Student’s t-tests were used. Results are presented as mean ± SD. * denotes a significant difference between pre– and post-training.

We distinguished three regimes of flows: Fåhræus-Lindqvist regime for vessel with a diameter above 8 µm, lubrication regime for vessels with a diameter below 7 µm and mixed regime for vessels with a diameter between 7 and 8 µm.

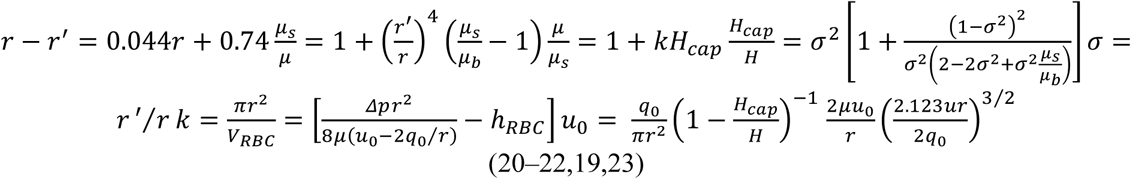

The network was then deformed to account for the pressure of blood inside the vessels. The structure of the walls is shown in Figure 2. For each generation, the final diameter was defined as the diameter providing a force balance between the pressure of the blood on the internal side and the elastic force of the walls and the intramuscular pressure on the external side.

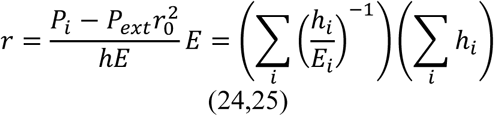

We assumed that the total flow in the muscles was 1 litre per minute at rest. For effort, we assumed that the increase in the cardiac output corresponded to an increase of the flow rate in the muscles and not in the other organs and adjusted the flow rate in the network accordingly. Finally, the angiogenesis was reproduced by doubling the number of capillaries while the rest of the vascular tree was left unaffected. The lengths and diameters of all the vessels corresponded to their pre-training dimensions to simulate a pure angiogenesis, without other remodelling. The length and diameter ratios between each vessel generation, a detailed explanation of the three flow regimes, and the elastic model of the vessels are provided in the Supplementary Methods (online appendix).

### Statistical analysis

Data normality was assessed for each variable using the Shapiro-Wilk test. Results are expressed as mean ± standard deviation. Paired Student’s t-tests were used to compare baseline and post-training (8 weeks) values. Two-way ANOVA was conducted with time and group as factors, and variance homogeneity was confirmed prior to analysis. Tukey’s post hoc test was applied when the ANOVA indicated a statistically significant effect. Statistical significance was accepted at p < 0.05.

## Results

A total of 30 young, healthy, and sedentary individuals were initially enrolled in the 8-weeks training intervention. Of the 30 participants, 25 successfully completed the intervention protocol (median age 21 years; 15 women and 10 men). Three participants (1 man and 2 women) failed to adhere to the training regimen, and two (2 women) sustained injuries unrelated to the exercise program. Finally, 13 performed MICT and 12 HIIT. No significant differences were observed in baseline characteristics between the groups (*Table 1*). All participants who completed the intervention demonstrated full adherence to the prescribed training protocol.

**Table 1.** Descriptive characteristics of participants.

| Characteristic | MICT<br>(n = 13) | HIIT<br>(n = 12) | p-value |
| --- | --- | --- | --- |
| Gender (F/M) | 9/4 | 6/6 |  |
| Age (years) | 21 (20 – 25) | 22 (20-27) | 0.722 |
| Weight (kg) | 70 ± 10 | 70 ± 11 | 0.896 |
| Height (cm) | 170 ± 12 | 171 ± 9 | 0.842 |
| BMI (kg/m<sup>2</sup>) | 24.5 ± 4.1 | 23.8 ± 2.2 | 0.595 |

BMI : body mass index. **Statistics :** Normality tests were performed by Shapiro-Wilk test. Student's t-test was applied for normally distributed data (mean ± SD) and Mann-Whitney U test for non-parametric data (median (25<sup>th</sup> -75<sup>th</sup> percentile).

### CPET

Aerobic capacity was similar between groups at baseline. As expected, VO₂_max_ increased in both groups following the training intervention, with a mean improvement of 4 ± 2 ml·kg⁻¹·min⁻¹ in the MICT group and 8 ± 5 ml·kg⁻¹·min⁻¹ in the HIIT group. The increase in VO₂_max_ was significantly greater in the HIIT group compared to MICT (p = 0.024). Consistently, maximal workload improved by 23 ± 13 watts in the MICT group and by 40 ± 18 watts in the HIIT group. Both VO₂ and workload at the first ventilatory threshold (VT1) showed significant improvements post-intervention in the MICT (respectively, p=0.032 and p=0.019) and HIIT groups (p<0.001 and p=0.007). CPET outcomes are summarized in *Table 2*.

**Table 2.** CPET data at before and following 8 weeks of training.

| CPET values | MICT (N=13) |  |  | HIIT (N=12) |  |  | Time* |
| --- | --- | --- | --- | --- | --- | --- | --- |
|  | baseline | 8 weeks | Time | Baseline | 8 weeks | Time | group |
| **Rest** |  |  |  |  |  |  |  |
| VO<sub>2</sub> (L. min<sup>-1</sup>) | 0.42 ± 0.07 | 0.44 ± 0.12 | 0.582 | 0.39 ± 0.08 | 0.42 ± 0.07 | 0.091 | 0.278 |
| VO<sub>2</sub> ( mL·kg<sup>-1</sup>·min<sup>-1</sup>) | 6.1 ± 0.9 | 6.4 ± 1.4 | 0.532 | 5.6 ± 1.1 | 5.9 ± 0.8 | 0.145 | 0.167 |
| HR (beat) | 93 ± 13 | 85 ± 13 | 0.082 | 84 ± 10 | 85 ± 13 | 0.743 | 0.113 |
| **VT1** |  |  |  |  |  |  |  |
| VO<sub>2</sub> (L. min<sup>-1</sup>) | 1.75 ± 0.35 | 1.94 ± 0.51 | 0.052 | 1.82 ± 0.44 | 2.23 ± 0.42 | **<0.001** | **0.038** |
| VO<sub>2</sub> (mL·kg<sup>-1</sup>·min<sup>-1</sup>) | 25 ± 4 | 28 ± 6 | **0.032** | 27 ± 6 | 31 ± 4 | **<0.001** | 0.538 |
| Workload (Watt) | 111 ± 27 | 129 ± 43 | **0.019** | 123 ± 43 | 149 ± 29 | **0.007** | 0.451 |
| HR (beat) | 148 ± 19 | 150 ± 13 | 0.467 | 137 ± 24 | 144 ± 20 | 0.279 | 0.526 |
| **Maximal** |  |  |  |  |  |  |  |
| VO<sub>2</sub> (L. min<sup>-1</sup>) | 2.61 ± 0.64 | 2.92 ± 0.75 | **0.006** | 2.56 ± 0.45 | 3.18 ± 0.75 | **<0.001** | **0.022** |
| VO<sub>2</sub> ( mL·kg<sup>-1</sup>·min<sup>-1</sup>) | 37 ± 7 | 41 ± 9 | **0.003** | 38 ± 6 | 46 ± 9 | **<0.001** | **0.024** |
| Workload (Watt) | 187 ± 49 | 210 ± 57 | **<0.001** | 198 ± 44 | 237 ± 51 | **<0.001** | **0.014** |
| HR (beat) | 184 ± 11 | 185 ± 11 | 0.567 | 177 ± 13 | 178 ± 16 | 0.585 | 0.978 |
| RER | 1.17 ± 0.07 | 1.16 ± 0.06 | 0.729 | 1.17 ± 0.08 | 1.16 ± 0.09 | 0.775 | 0.862 |

VO<sub>2</sub> : Oxygen uptake, HR : Heart rate, VT1 : first ventilatory threshold, RER : Respiratory Exchange Ratio, Baseline : before 8-weeks training, 8 weeks : After 8-weeks training. **Statistics** : Normality tests were tested with a Shapiro-Wilk test. For pairwise comparisons, Student's t-tests were used. ANOVA tests were performed to compare groups and Tukey post hoc tests were applied. Results are presented as mean ± SD.

### Non invasive determination of Cardiac output

Out of the 25 participants, 15 performed non-invasive Q measurements (8 assigned to MICT and 7 to HIIT). At peak exercise, Q_max_ increased similarly in both groups after 8 weeks of exercise training: from 16 ± 3 L/min to 19 ± 4 L/min (+ 13 ± 7%, p=0.002) in MICT and from 18 ± 3 L/min to 21 ± 2 L/min (+ 15 ± 9%, p=0.001) in HIIT. In parallel, stroke volume increased from 88 ± 13 to 101 ± 17mL (+ 15±8 %, p=0.01) in the MICT group and from 100 ± 15 to 119 ± 16mL (+ 19 ± 13%, p<0.001) in the HIIT group. No modification of maximal heart rate was observed. Within this specific subgroup, VO₂_max_ also increased, with a greater improvement observed in the HIIT group compared to MICT (p = 0.049). Calculated peripheral O₂ extraction remained unchanged in the MICT group (16±3 to 16±2 mlO2/100ml, p=0.984), while a tendency toward an increase was observed in the HIIT group (14±2 to 16±3 mlO2/100ml, p=0.066). However, a significant group-by-time interaction was detected (p = 0.041), suggesting a differential adaptation in peripheral oxygen extraction between training modalities. Within the whole group, VO₂_max_ correlated with Q_max_ before and after 8 weeks of training, respectively: r=0.630, p=0.012 and r=0.847,p<0.001.

### Resting muscle biopsies outcomes

In a subgroup of 10 participants equally divided between MICT (n=5) and HIIT (n=5), arteriolar intima diameters exhibited non-significant enlargement from 20 ± 3 μm at baseline to 26 ± 4 μm post-training in the MICT group (p=0.054), and from 22 ± 5 μm to 27 ± 5 μm in the HIIT group (p=0.171). Similarly, media diameters of arterioles showed a comparable dynamic, from 30 ± 4 μm to 35 ± 7 μm in MICT (p=0.181), and from 31 ± 9 μm to 41 ± 12 μm in HIIT (p=0.147). Capillary intima diameters increased following training, from 6.2 ± 0.6 μm to 7.2 ± 0.5 μm in MICT (p = 0.001) and from 6.3 ± 0.7 μm to 7.4 ± 0.5 μm in HIIT (p = 0.045). The media diameters of capillaries (representing pericytes coverage) showed no significative increase, from 7.1 ± 0.6 μm to 7.7 ± 0.5 μm in MICT (p = 0.06), and an eventual increase from 7.3 ± 0.6 μm to 8.3 ± 0.5 μm in HIIT (p = 0.047) (Figure 4, E and J). No pro-angiogenic response was detected. Three-dimensional analyses revealed no detectable expansion of the capillary network across conditions. In the MICT group, mean capillary density was 3135 capillaries/mm³ at baseline and 2931 capillaries/mm³ post-training (p=0.231). Similarly, the HIIT group exhibited values of 3131 capillaries/mm³ at baseline and 3086 capillaries/mm³ following the intervention (p=0.805) (Figure 4, D and I). Statistical analysis confirmed the absence of significant differences over time*groups (p = 0.499). Venular internal diameters demonstrated a pronounced increase, from 28 ± 8 μm to 38 ± 10 μm in MICT (p = 0.015), and from 23 ± 6 μm to 36 ± 6 μm in HIIT (p = 0.003). Correspondingly, venular external diameters expanded, from 36 ± 11 μm to 45 ± 13 μm in MICT (p = 0.009), and from 32 ± 10 μm to 44 ± 7 μm in HIIT (p = 0.021) (Figure 4, G and L).

### Hemodynamic model

Based on estimation from our vascular model *(Figure 5, panel A)*, capillary shear stress decreased in response to both training modalities after eight weeks. Under training-intensity conditions (i.e. 65% HR_max_ for MICT and 90% HR_max_ for HIIT), capillary shear stress declined from baseline to post training intervention: from 7.45 Pa to 6.16 Pa in MICT and from 18.15 Pa to 10.47 Pa in HIIT. These values indicate that baseline shear stress (i.e., before training) was higher than post-training shear stress under the same intensity conditions (*Figure 5, panel B)*. The ratio between the relative reduction in shear stress during training and the increase in pericyte coverage yielded similar values for both modalities: 17.6 MPa·m⁻¹ for MICT and 18.35 MPa·m⁻¹ for HIIT *(Figure 5, panel C).* Similarly to capillaries, venular shear stress also decreased from baseline to post-training. Under the same intensity conditions, venular shear stress dropped from 9.55 Pa to 4.69 Pa with MICT and from 26.90 Pa to 8.79 Pa with HIIT (figure 5, panel D). The ratios between changes in venular shear stress and corresponding alterations in venular diameter was consistent: 3.66 for MICT and 3.80 for HIIT (Figure 5, panel E). These ratios reflect the relationship between the reduction in venular shear stress during training and the corresponding increase in venular diameter. Furthermore, as illustrated in *Figure 5, panel F,* we hypothesize that the enhancement of capillary pericyte coverage under HIIT conditions may contribute to mechanical stabilization of the capillary wall, potentially limiting deformation under elevated shear forces. Finally, to evaluate the potential impact of capillary angiogenic expansion, we simulated pressure drops across two vascular architectures *(Figure 5, panel G)*: one comprising five capillary generations, which corresponds to our model, and another incorporating a hypothetical sixth generation to mimic angiogenic development. The five-generation model yielded an average pressure drop of 85.97 mmHg, while the six-generation model produced a pressure drop of 200.66 mmHg, nearly double the initial value.

**Figure 5:**
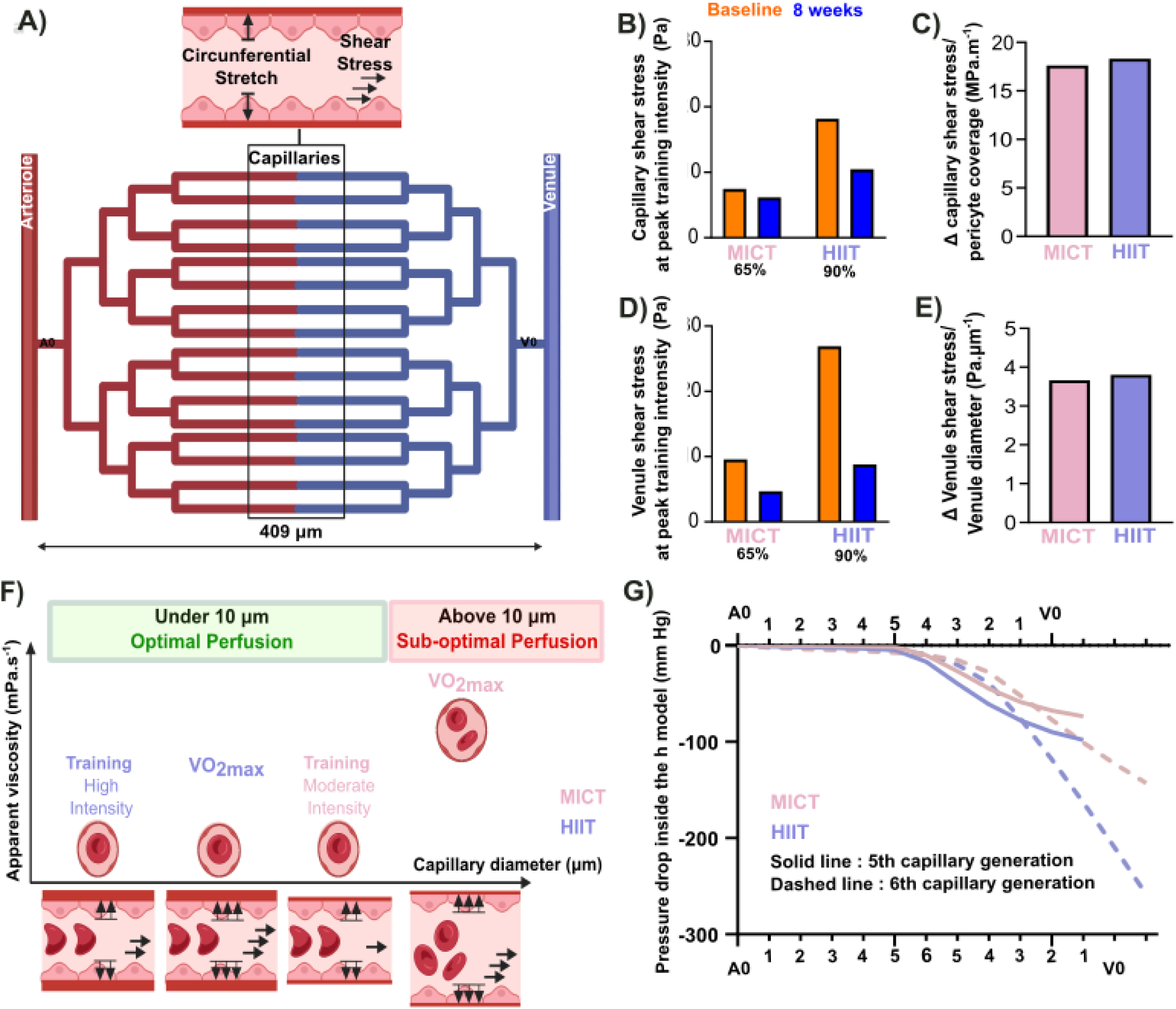
Hemodynamic model and derived measurements. (A) depicts the microvascular hemodynamic model developed to simulate flow dynamics across a branching capillary network. The structure consists of a single arteriole outlet (AO) bifurcating into five successive generations of capillaries, culminating in 32 individual capillary segments that reconverge into a single venule outlet (VO). The total length of the model spans 409 μm. Hemodynamic forces were estimated throughout the network under physiologically relevant conditions. (B – E) show all shear stress values at peak training intensity (i.e. 65% HR_max_ for MICT and 90% HR_max_ for HIIT). Panel B displays capillary shear stress in both training group. (C) shows the ratio between training-induced changes in capillary shear stress and the relative variation in pericyte coverage following the 8-week intervention. (D) reports venular shear stress values across MICT and HIIT conditions. (E) illustrates the relationship between changes in venular shear stress and corresponding alterations in venular diameter. (F) characterizes apparent blood viscosity as a function of vessel diameter, highlighting red blood cell (RBC) morphology and axial positioning in capillaries below and above 10 μm in diameter. The number and orientation of flow vectors reflect the magnitude and directionality of local hemodynamic forces within the capillary bed. (G) compares pressure drop profiles at VO₂max in models containing five versus six capillary generations under both MICT and HIIT conditions.

## Discussion

Our results indicate a specific and unexpected pattern of capillary adaptation modulated by training intensity. While no capillary angiogenesis was observed, capillary diameter expansion appears to be constrained by pericyte proliferation. Our hemodynamic model predicts that this adaptation contribute to lumen optimization to improve centring of RBC convection and facilitate oxygen diffusion within muscle fibers. An improved oxygen extraction and utilization during elevated levels of cardiac output would therefore facilitate aerobic exercise. We further observed that venular remodelling followed the principles of the shear stress set-point theory, where blood vessel diameter adapts in response to changes in shear stress, to restore the original shear stress set point(11,14).

### Aerobic capacity

Our results demonstrate that both MICT and HIIT induce significant improvements in VO_2max_ over an eight-week intervention in sedentary young adults. In parallel, the magnitude of improvement in VO₂_max_ was significantly greater following HIIT, reinforcing the growing body of evidence supporting its superior efficacy in enhancing aerobic capacity (4,26). This differential effect appears to be particularly pronounced in healthy individuals aged 14 to 45 years (5) and influenced by baseline fitness level, while remaining largely independent of sex (7). These effects were also observed by Milanović et al. (6) reporting greater improvement in VO₂_max_ following HIIT, compared to MICT under matched training volumes, suggesting that training intensity is a key driver of aerobic adaptations. It is noteworthy that the magnitude of VO₂_max_ improvement observed in our HIIT group exceeds that reported in the meta-analysis of Scribbans et al (4) that found a mean increase in VO₂_max_ of 4.9 ± 1.4 ml·kg⁻¹·min⁻¹ for MICT versus 5.5 ± 1.2 ml·kg⁻¹·min⁻¹ for HIIT what could partly be explained by the substantial heterogeneity in HIIT protocols across studies. Manipulation of the FITT principle which refers to frequency, intensity, time and type, has indeed emerged as a critical determinant of both acute and chronic, physiological and metabolic responses to interval training (27). Wen et al. (28) accordingly reported that protocols incorporating intervals of at least 2 minutes in duration, combined with a cumulative high-intensity workload of no less than 15 minutes per session over a period of 4 to 12 weeks, consistently produced greater improvements in aerobic fitness compared with MICT. Similarly, Guo et al. (5) showed also that training with high intensity intervals lasting between 1 and 3 minutes, performed three times per week for a minimum of six weeks, optimize cardiorespiratory adaptations. The superior efficacy of longer HIIT bouts likely stems from sustained near-maximal intensity, enhancing central and peripheral cardiovascular stimuli (8). The greater VO₂_max_ improvement observed in our study may reflect our protocol’s specific design of 14 minutes cumulative training of 2 minutes intervals at 90% of HR_max_.

### Peripheral oxygen extraction

In our subgroup of 15 participants VO₂_max_ improved across both training modalities with a parallel and similar increase in Q_max_. These observations are in line with VO_2max_ increase being primarily related to an increase in maximal oxygen transport (i.e. Q_max_) to the exercising muscle, as described in several meta-analyses (29,30). However, compared to the MICT group, VO_2max_ increased slightly more in the HIIT group, suggesting a better oxygen extraction at maximal exercise. Although the arteriovenous oxygen difference (a-vO_2_ diff) remained generally stable among our participants, we observed a non-significant trend toward a greater increase in the HIIT group (14±2 to 16±3 mlO2/100ml, p=0.66) that was accompanied by a temporally divergent trajectory between training modalities (p=0.041). Oxygen extraction is considered to result from a combination of factors, including an intensified Bohr effect (31), prolonged erythrocyte transit time, increased capillary density (32), optimized vascular recruitment and flow distribution (33), and greater oxidative capacity of muscle tissue (34). In that context, greater exercise intensity during HIIT appears to amplify muscle intracellular signaling cascades, resulting in a superior increase in mitochondrial content compared to work-match MICT protocol (35). Moreover, repeated exposure to elevated hemodynamic forces, which characterizes HIIT, appears to induce superior vascular adaptations compared to traditional endurance training. (36). Gene and protein expression analyses, from human vastus lateralis of the quadriceps, demonstrate indeed that low-volume, high-intensity exercise transcriptionally activates angiogenic factors (2) suggesting promotion of better organised vasculature for exercise oxygen disponibility. However, the extent to which aerobic training procedure specificity enhances muscle oxygen extraction remains uncertain. A meta-analysis conducted by Montero et al. (37) reported no significant improvement in a-vO₂ diff following aerobic training (mixing MICT and HIIT protocols), although studies with longer duration or higher load of training induced a higher a-vO2 diff. However, as highlighted by Poole (38), these reports should be interpreted with caution as of some of the studies included in this meta-analysis were subject to methodological limitations related to a limited use of catheter-based measurements and potential inaccuracies in VO₂max and Qmax estimations. In contrast, a more recent meta-analysis by Astorino et al. (30) suggests that aerobic training, particularly when involving high-intensity intervals, can elicit modest but significant increases in a-vO₂ diff. Finally, our results tend to confirm HIIT superiority in improving a-vO2 diff at maximal exercise that may stem from vascular remodelling processes, which could facilitate more efficient oxygen extraction as explained below.

### Vascular remodelling

While macrovascular adaptations to exercise have been extensively documented, data regarding microcirculatory remodelling in humans remain limited. Thijssen et al. (39) demonstrated a dose-dependent improvement in femoral artery flow-mediated dilation (FMD) and reduced arterial wall thickness following aerobic training. Meta-analyses further support a positive correlation between exercise intensity and endothelial function (40), with HIIT yielding greater macrovascular benefits than MICT (41). In contrast, little to no data exist describing structural adaptations at the microvascular level (arteriolar, capillary, and venular diameters), as much of the current understanding stems from animal models, and in vivo technical challenges restrict investigation within human quadriceps muscle. To our knowledge, our study is the first to assess microcirculation diameters and angiogenic adaptations in human skeletal muscle using advanced tissue clearing techniques combined with high-resolution confocal microscopy, which enables the three-dimensional visualization of entire human muscle samples, along with micron-level precision for measuring diameters from microcirculation and conducting longitudinal capillary quantification. Direct comparison with previous literature is therefore limited, as most studies on exercise-induced angiogenesis have relied on conventional two-dimensional histological sections with transverse slices stained with immunomarkers to estimate capillary density (capillaries per mm²) or capillary-to-fiber ratios. Other investigations have focused on the expression of angiogenesis-related biomarkers, especially the vascular endothelial growth factor (VEGF) (12,42). From those studies, capillary density is estimated to range between ∼300 and 600 capillaries per mm², depending on training status (32) with potential increase of 49.7 capillaries/mm² following endurance training (15% improvement), and that exercise intensity is a key modulator of an angiogenetic response (43). Interestingly, numerous studies (44) (45) (10) have identified shear stress as a key biomechanical stimulus for angiogenesis. Despite extensive literature supporting exercise-induced angiogenic responses in human skeletal muscle, our study did not demonstrate clear evidence of new capillary formation, which methodological and physiological factors could explain. Indeed, the use of advanced tissue clearing combined with high-resolution confocal microscopy and immunostaining allowed for longitudinal visualization of the vascular network with micron-level precision that may offer superior anatomical fidelity to enable identification of three-dimensional vessel continuity and connectivity. A two-dimensional transverse section might erroneously count capillaries due to vessel bifurcation or anastomosis, thereby potentially influencing the estimate of angiogenesis in previous studies. Moreover, we observed a general increase in capillary diameters after training, suggesting pre-existing capillaries dilatation rather than the formation of entirely new ones. The increased vascular caliber likely improves the detectability of pre-existing capillaries that existed before training but fell below the resolution threshold. In this context, structural changes may reflect vascular remodelling rather than de novo angiogenic activity. Further, our hemodynamic model demonstrates that adding another generation of capillaries in a branched network results in huge pressure drops, which would hinder the optimal perfusion of the muscle.

### Modelisation and interpretation

In humans at rest, typical shear stress values range between 0.5 and 5 Pa (11). Numerous studies (15,36,46) have introduced the concept of a “set point”, whereby endothelial cells drive vascular remodelling, adapting vessel diameter and luminal area to maintain shear stress within an optimal physiological range. A particularly compelling insight from the work of Baeyens et al. (11) is that the set point may differ across vessel types. In their study, comparison of endothelial cells from veins and lymphatic vessels highlighted the functional heterogeneity of endothelial responses. In our exercise-based model, structural adaptation following blood flow elevation and increased shear stress stimulated both capillaries and venules. Since the biopsies were obtained at rest, the adaptations observed in the capillary and venular segments represent permanent structural changes, in addition to functional vessel adaptation. However, a more pronounced arteriolar vasoreactivity, as evidenced by improved FMD in response to training (40), might still exist without structural modifications.

Furthermore, our study demonstrated a significant increase in pericyte coverage at the capillary level following HIIT, whereas MICT did not produce a similar effect. This divergent pattern suggests that HIIT may trigger the activation of protective mechanisms to regulate microvascular architecture. We hypothesize that the increased pericyte coverage observed under HIIT may help stabilize the microvessel structure in response to high mechanical stimuli, thereby preserving an optimal diameter for oxygen exchange *(Figure 4, panel G)*. From a rheological standpoint, vessel diameter has profound effects on blood viscosity and the spatial dynamics of red blood cells (RBCs) (47). In vessels larger than 10 μm, multiple RBCs can travel side-by-side, interact and shift away from the vessel center, leading to peripheral displacement and increased apparent viscosity. In small vessels (diameter <10 μm), red blood cells (6–8 μm) deform and align in single file, moving centrally with a thin plasma layer separating them from the vessel wall, a behavior described by the Fåhræus–Lindqvist effect. This microvascular flow configuration is especially favourable for oxygen diffusion, as it increases the surface area available for RBC gas exchange making microvascular oxygen transport tightly related to capillary geometry and RBC alignment. Studies have shown indeed that when RBCs travel centrally and in single file, oxygen extraction becomes more efficient due to minimized diffusion barriers (47). We propose that the increased pericyte coverage observed in HIIT may help stabilize capillary diameter against pressure-induced deformation, thereby preserving this optimal flow configuration. This interpretation is further supported by our post-training analysis of the shear stress-to-pericyte coverage ratio yielded comparable ratios in both protocols inducing that pericyte coverage increases proportionally to the shear stress experienced during training. Such adaptation likely acts as a biomechanical safeguard, limiting excessive capillary deformation and helping maintain an optimal vessel diameter for oxygen exchange at specific training intensities. When extrapolated to maximal effort, this mechanism may carry important functional implications as when blood flow and shear stress surge, vessels conditioned through MICT, (i.e. lower shear stress) will exceed their tolerance, leading to over-dilation and reduced oxygen diffusing ability. In contrast, HIIT-trained vessels, conditioned to higher shear stress, are better equipped to withstand peak hemodynamic loads and preserve optimal flow geometry at maximal exercise intensity. Interestingly, veins adapted in accordance to the set point theory (14) and consistent with a homeostatic response to shear stress elevation, modulating luminal diameter to stabilize local hemodynamic stress. Indeed diameter increase following training was proportionate to hemodynamic forces increases in MICT or HIIT (Fig. 5, panel D and E). To note, Tiezzi (48) et al. mentioned shear stresses in veins ranging from 0.1 to 0.6 Pa in healthy individuals. In our cohort, resting shear stress levels decreased after training from 0.9 to 0.7 Pa in the MICT group and from 1.3 to 0.8 Pa in the HIIT group. These higher values are probably explained by the smaller diameters of venules compared to veins. Finally, no evidence of angiogenesis was observed following exercise training. This contradicts the current paradigm, as exercise is generally associated with the induction of angiogenesis. However, simulating an additional generation of capillaries in our prediction model resulted in a pressure drop across the remodelled network of approximately 200.66 mmHg, producing flow conditions that are incompatible with microvascular viability. In this configuration, the addition of a capillary generation rendered the system mechanically unstable and unable to sustain coherent perfusion. Our article therefore proposes that microvascular adaptation to aerobic exercise is mainly dictated by mechanotransduction rather than chemically-induced angiogenesis.

### Limitation

Vesel diameters were measured at rest, as it is impossible to fix perfused biopsies. Second, blood samples were not collected, which prevented a direct assessment of haemoglobin concentration. Given haemoglobin’s influence on blood viscosity and oxygen-carrying capacity, its absence limits the precision of rheological estimates. Third, cardiac output measurements and muscle biopsy data were collected from two distinct subject subgroups. Although both populations were matched for age, training status, and cardiovascular fitness, the absence of direct overlap introduces a methodological constraint. Consequently, while the findings offer valuable insight into microvessel behavior, they may lack the granularity required to establish individual-level associations between flow dynamics and vascular remodelling.

In conclusion, our study demonstrates that high-intensity interval training elicits a more pronounced increase in VO₂_max_ compared to moderate-intensity continuous training, despite both protocols inducing similar enhancements in maximal cardiac output. We observed training intensity-dependent remodelling in pericyte capillaries, with no evidence of angiogenesis, to maintain optimal diameter and optimize muscle perfusion and oxygen extraction under the highest metabolic hemodynamic stress.

## Acknowledgements

We express our deep gratitude to all study participants for their time and commitment.

## Source of funding

No financial support, including funds or grants, was provided to the authors during the writing of this manuscript. The confocal microscope used in this study was acquired thanks to funding of the Fondation ULB.

## Disclosures

None

## Notes

### Competing Interest Statement

The authors have declared no competing interest.

